# A temporally dynamic *Foxp3* autoregulatory transcriptional circuit controls the effector Treg programme

**DOI:** 10.1101/238386

**Authors:** David Bending, Alina Paduraru, Paz Prieto Martin, Tessa Crompton, Masahiro Ono

## Abstract

Regulatory T cells (Treg) are negative regulators of the immune response. Whilst thymic Treg generation is well studied, it is not known whether and how *Foxp3* transcription is induced and regulated in the periphery during immune responses. Here we use Foxp3 **T**imer **o**f **c**ell **k**inetics and activit**y** (Tocky) mice, which report real-time *Foxp3* gene transcription by measuring the spontaneous maturation of Fluorescent Timer protein from Blue to Red fluorescence, to identify the flux of *Foxp3*-to *Foxp3*+ T cells within the periphery and analyse the real-time activity of *Foxp3* transcription. Using a murine model of skin allergy, we show that both the flux of new *Foxp3* expressors and the rate of *Foxp3* transcription are increased at inflamed sites. These persistent dynamics of *Foxp3* transcription determine the effector Treg programme, and are dependent on a Foxp3 autoregulatory transcriptional circuit, as evidenced by analysis of T cells lacking functional Foxp3 protein. Such reactive and persistent *Foxp3* transcriptional activity controls the expression of coinhibitory molecules including CTLA-4 and effector-Treg signature genes. Using RNA-seq, we identify two groups of surface proteins based on their relationship to the temporal dynamics of *Foxp3* transcription, and we show proof-of-principle for the manipulation of *Foxp3* dynamics by immunotherapy: new *Foxp3* flux is promoted by anti-TNFRII antibody, and high frequency *Foxp3* expressors are depleted by anti-OX40 antibody. Collectively, our study dissects time-dependent mechanisms behind Foxp3-driven T cell regulation, and establishes the Foxp3-Tocky system as a tool to investigate the mechanisms behind T cell immunotherapies.

## Introduction

Upon antigen recognition through the T cell receptor (TCR), T cells express Interleukin(IL)-2 and CD25 (IL-2 receptor alpha chain), which together promote T cell activation, proliferation, and differentiation (1, 2). Intriguingly, CD25-expressing T cells from healthy animals are markedly enriched with regulatory T cells (Treg) that express the transcription factor Foxp3 (3, 4). Foxp3 expression is a major determinant of Treg phenotype and function, and Foxp3 interacts with transcription factor complexes, such as those involving NFAT and Runx1, to repress IL-2 transcription and convert the effector mechanisms in T cells into a suppressive one (5-7). Treg have activated phenotypes, and upon TCR signals, Treg suppress the activities of conventional T cells (8). TCR signaling is the major regulator of Treg differentiation in the thymus, as T cells that have received strong TCR signals preferentially express CD25 and Foxp3 and differentiate into Treg (9). Additionally, costimulatory receptors augment TCR signal-dependent Foxp3 and CD25 expression (10, 11). In the periphery, strong TCR signals further differentiate Treg into “effector Treg”, showing enhanced suppressive function (12).

Accumulating evidence indicates that Foxp3 expression is dynamically controlled in Treg and non-Treg. TCR stimulation induces Foxp3 expression in human (13) and mouse T cells (14) *in vitro*. Studies using T-cell receptor (TCR) transgenic systems have shown that Foxp3 expression is induced in non-Treg in some inflammatory conditions in vivo (15). Although such induced Foxp3 expression is often dismissed as “transient expression”, the dynamic induction of Foxp3 expression may have functional roles during T cell responses if this reactive Foxp3 expression occurs in activated polyclonal T cells during inflammation *in vivo*(16). In addition, Foxp3 expression can be dynamically downregulated in Treg. Fate-mapping experiments showed that, while most of thymus-derived Foxp3+ T cells stably express Foxp3, some Foxp3+ cells downregulate Foxp3 to become ex-Foxp3 cells in the periphery, joining the memory-phenotype T cell pool (14). PD-1 KO mice with a partial Foxp3 insufficiency leads to generation of ex-Foxp3 effector T cells (17), indicating that the mechanism of T cell activation is involved in the dynamic regulation of *Foxp3* transcription. These findings lead to the hypothesis that Foxp3 acts as a cell-intrinsic and transcellular negative feedback regulator for T cell activation among self-reactive T cell repertoires (16), challenging the thymus-central view of Treg-mediated immune regulation.

The key question is whether and how frequently activation of new *Foxp3* transcription is induced in non-Treg cells in physiological conditions, and how *Foxp3* transcription is sustained in existing Treg during the immune response. Since the death rate of Treg and other T cells is difficult to determine experimentally, the relative proportions of Foxp3+ and Foxp3-cells in steady-state conditions may not reflect the probability of new *Foxp3* induction in individual T cells, especially when T cells are expanding and dying during the immune response. Furthermore, human studies show that the level of Foxp3 expression may determine the functional state of Treg: the higher Foxp3 expression is, the more suppressive Treg are (18, 19). Thus, it is fundamental to investigate the temporal dynamics of *Foxp3* transcription over time in individual T cells in vivo, but this has been technically difficult to do to date.

Here we use our novel **T**imer **o**f **C**ell **K**inetics and Activit**y** (Tocky) system (20) to reveal the time and frequency of *Foxp3* transcription during peripheral immune responses. In the Foxp3-Tocky system, the transcriptional activity of the *Foxp3* gene is reported by Fluorescent Timer protein, which emission spectrum spontaneously changes from Blue to Red fluorescence after translation (21), and the in vivo dynamics of *Foxp3* transcription is analysed. We show that during inflammation, the flux of *Foxp3*-to *Foxp3*+ T cells dramatically increases within the periphery. In addition, we demonstrate that the real-time frequency of *Foxp3* transcription determines the effector Treg differentiation. Thus, we provide experimental evidence that *Foxp3* expression is dynamically regulated in Treg and non-Treg during inflammation in vivo, providing fresh insight into Foxp3-driven T cell regulation.

## Results

### Timer-Blue fluorescence reports real-time *Foxp3* transcription

Fluorescent Timer protein (Timer) is an mCherry mutant, and when translated, the chromophore of Timer is an unstable blue-form, which spontaneously and irreversibly matures to become a stable red-form (21). We hypothesised that Timer-Blue fluorescence in Foxp3-Tocky mice reports the real-time transcriptional activities of the *Foxp3* gene. To determine the relationships between *Timer* mRNA expression and endogenous *Foxp3* transcripts, we performed an RNA degradation assay using actinomycin D. After actinomycin D treatment, the transcripts of *Timer*, *Foxp3*, and an unrelated mRNA species, *Hprt*, exhibited similar half-lives (1.14 h, 1.46 h, and 1.73 h, respectively, **Fig. 1A**). This indicates that *Timer* transcripts are well correlated to *Foxp3* ones in Foxp3-Tocky T cells, in which *Timer* transcripts report the transcriptional activity of the *Foxp3* gene (20). Next, we analysed the half-life of Timer-Blue fluorescence in activated T cells *in vitro* using a short-term treatment with cycloheximide (CHX) to inhibit new protein synthesis. Whilst a previous study estimated the maturation half-life of Timer-Blue to be 7.1 h, using purified Timer proteins and by fitting data to a pharmacological kinetic model (21), here we aimed to experimentally determine the half-life of Timer-Blue fluorescence by flow cytometric analysis of Foxp3-Tocky T cells. Upon CHX treatment, flow cytometric plots showed that Foxp3-Timer Blue fluorescence rapidly decayed, while Timer-Red fluorescence was more stable (**Fig. 1B**). The half-life of Timer-Blue fluorescence was estimated to be 4.1 h (**Fig. 1C**) whilst Timer-Red fluorescence gradually increased (**Fig. 1D**), due to the accumulation of matured proteins. The short half-life of Timer-Blue fluorescence indicates that Timer-Blue fluorescence reports real-time *Foxp3* transcripts, while Timer-Red fluorescence captures the cumulative activity of *Foxp3* transcription over time.

**Figure 1:**
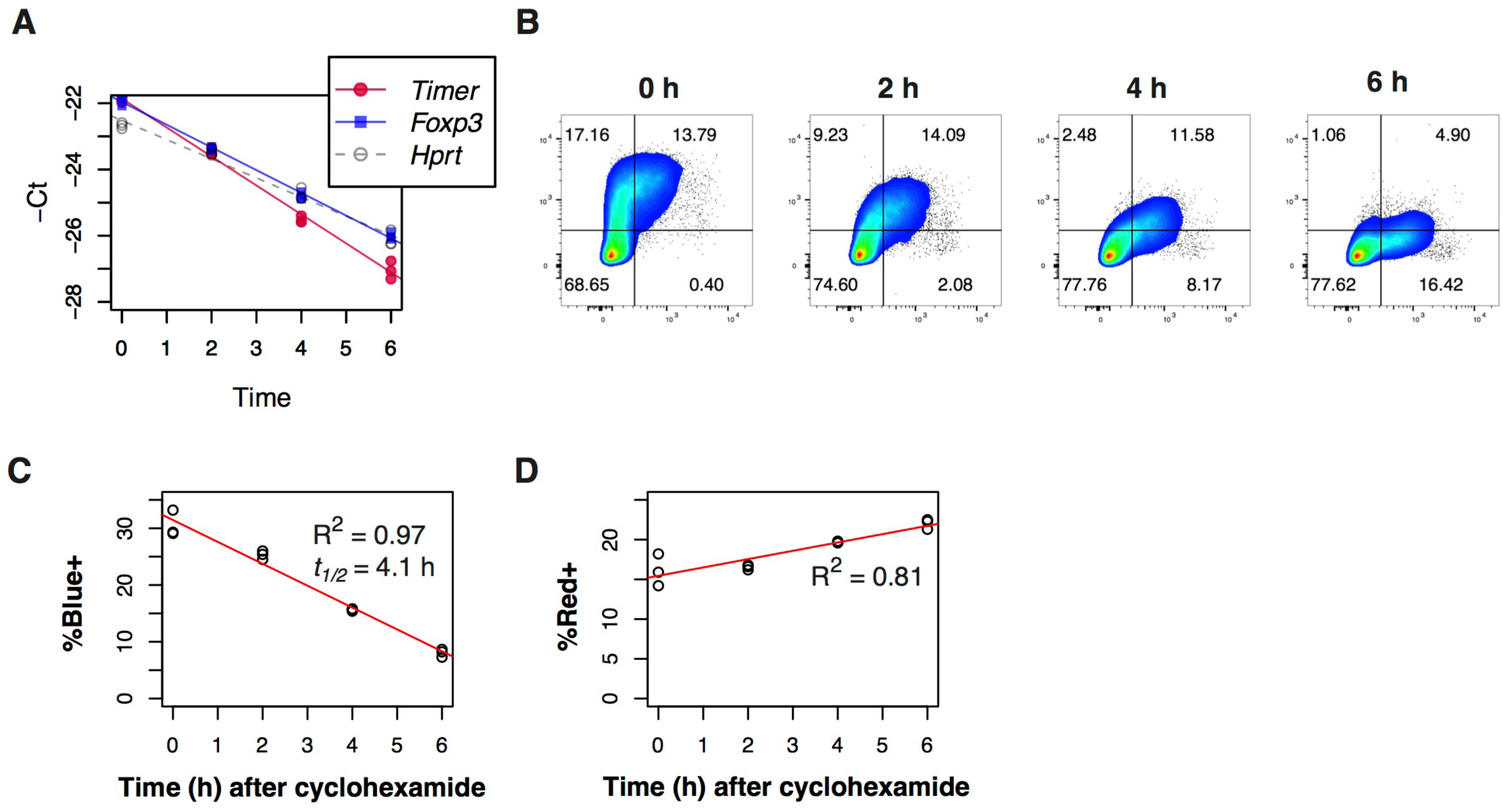
Timer-Blue fluorescence reports real-time *Foxp3* transcription. **(A)** CD4+ T cells from Foxp3-Tocky mice were cultured for the indicated time points with 10μg/ml actinomycin D. RNA was extracted and *Timer*, *Foxp3* and *Hprt* mRNA detected by RT-PCR. Plotted are the raw Ct values. (**B**) Naïve T cells from Foxp3-Tocky mice were stimulated for 48hrs in the presence of anti-CD3 and anti-CD28 in the presence of IL-2 and TGFb. After 48hrs cells were harvested and incubated with 100μg/ml cycloheximide for the indicated time points. Cells were analysed by flow cytometry, shown are the Timer-Blue vs. Timer-Red fluorescence within CD4^+^ T cells. Summary data of % Timer-Blue+ (**C**) or % Timer-red+ (**D**) in the cultures following addition of cycloheximide. Linear regression analysis by Pearson’s correlation. Data are representative of two independent experiments.

### Thymic CD4-single positive cells have higher rates of new *Foxp3* transcription compared to splenic CD4+ T cells in neonatal mice

In neonatal mice, Foxp3+ T cells are actively produced in the thymus (22). In Foxp3-Tocky mice, thymic CD4-single positive cells showed a higher percentage of Blue+ cells in Timer+ cells than splenic CD4+ cells (i.e. 95% and 63% of all Timer+ cells were Blue+, **Fig. 2A**). Timer-Blue expression levels in Timer+ cells were much higher in the thymus than in the spleen, while Timer-Red expression levels were higher in the spleen (**Fig. 2B**). These data indicate that the rate of *Foxp3* transcription is higher in the thymus than the spleen, while splenic Foxp3+ cells have transcribed the *Foxp3* gene for a longer time on average than thymic Foxp3+ cells. These results thus further confirm that Foxp3-Tocky captures real-time *Foxp3* transcription by Timer-Blue fluorescence and its history and cumulative activity by Timer-Red in vivo.

**Figure 2:**
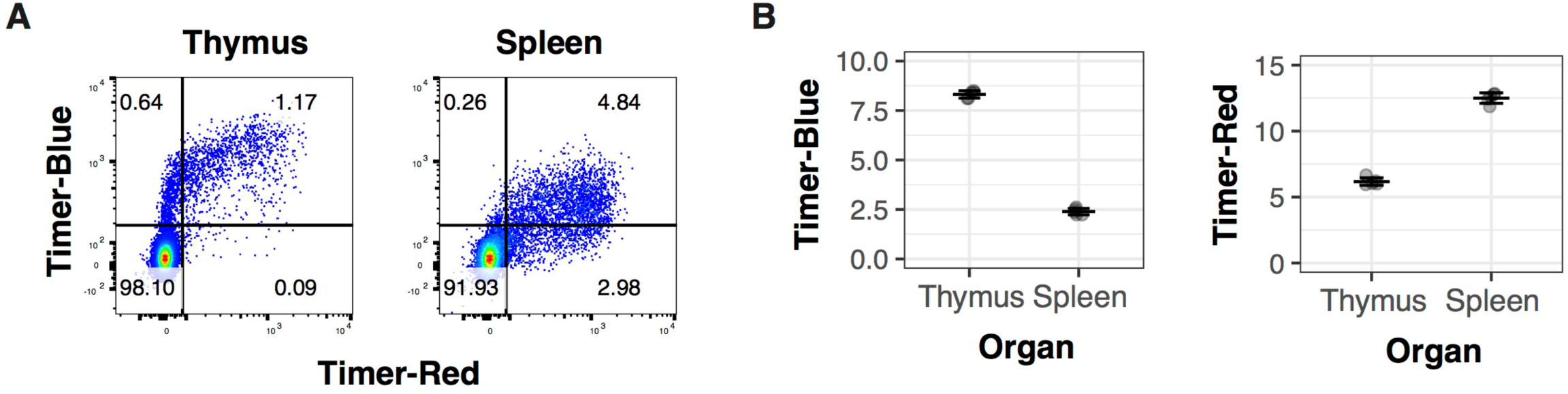
Thymic CD4-single positive cells have higher rates of new *Foxp3* transcription compared to splenic CD4+ T cells in neonatal mice. (**A**) CD4-single positive cells from the thymus and CD4^+^ T cells from the spleens of day 10 old Foxp3-Tocky mice were analysed for Timer-Blue vs. Timer-Red expression by flow cytometry. (**B**) Summary data of Timer-Blue and Timer-Red MFI of CD4^+^ T cells from thymnus or spleen of individual Foxp3-Tocky mice, n=5.

### The flux of new *Foxp3* expressors and the rate of *Foxp3* transcription are increased in tissue-infiltrating T cells during the immune response

Next, we tested if *Foxp3* transcriptional activity is dynamically regulated during the immune response. We used a skin contact hypersensitivity (CHS) model to analyse the dynamics of *Foxp3* transcription in draining lymph nodes (dLN) and skin-infiltrating T cells after sensitising mice with the hapten oxazolone and challenging them on their ears (**Fig. 3A**). T cells were not present in sufficient numbers to be recovered from Vehicle-treated control skins. Most Timer+ cells in dLN were Red+, and these expressed various levels of Blue fluorescence (**Fig. 3B**), which indicates that *Foxp3* transcription levels vary between individual cells. While immunisation with oxazolone resulted in minimal effects in dLN, skin-infiltrating T cells showed uniformly high Timer-Blue fluorescence, indicating that these cells had high-frequency persistent *Foxp3* transcription (**Fig. 3B and 3C**). On the other hand, Timer-Red fluorescence was lower in skin-infiltrating T cells than dLN cells, indicating that these skin T cells had less accumulated *Foxp3* transcripts, and thus that these cells are short-lived or rapidly dividing.

**Figure 3:**
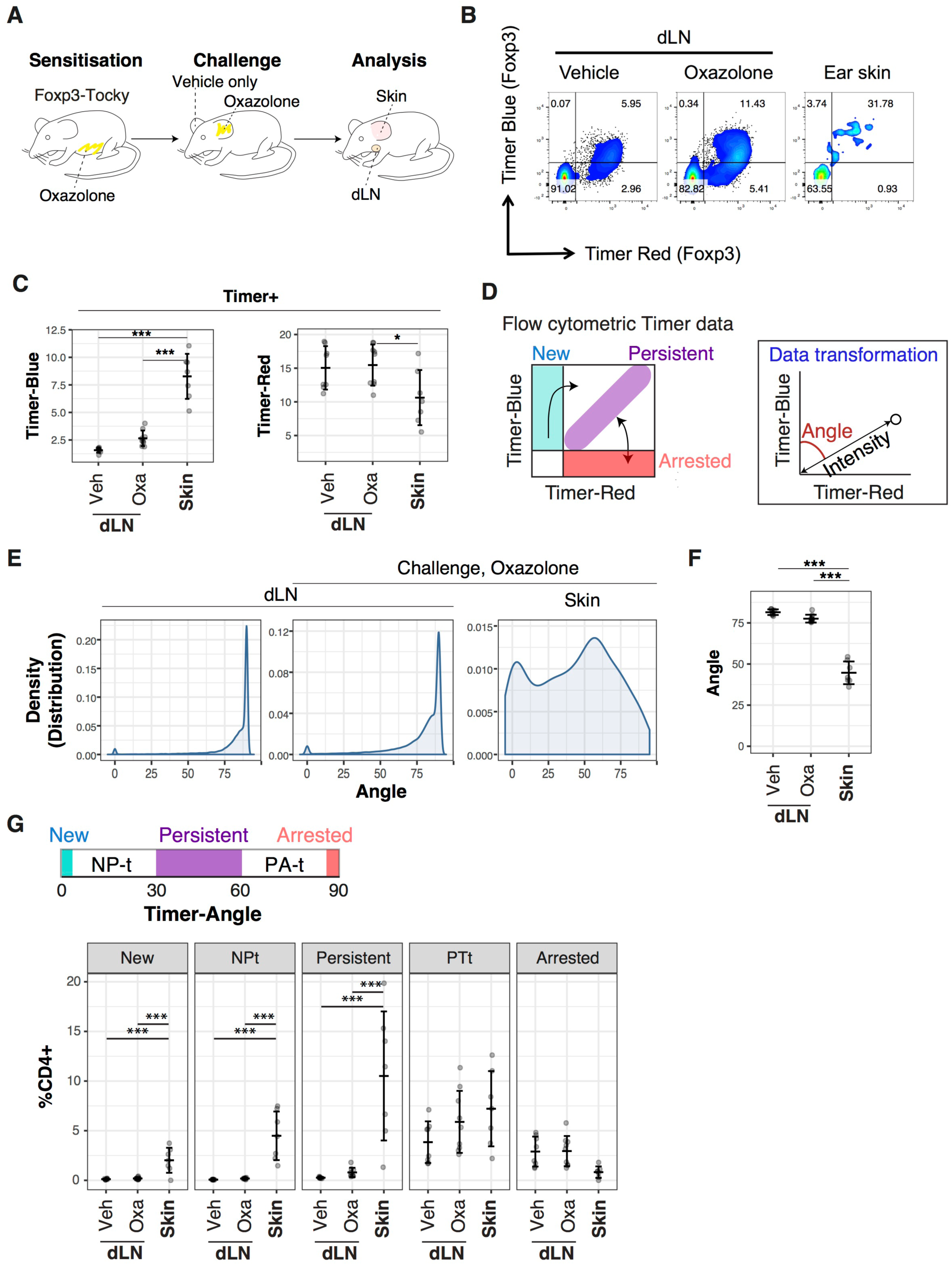
The flux of new *Foxp3* expressors and the rate of *Foxp3* transcription are increased in tissue-infiltrating T cells during the immune response. (**A**) Schematic displaying experimental set-up for analysis of sensitisation and challenge phases of Oxazolone-induced CHS using *Foxp3-Tocky mice*. (**B**) 96hrs after challenge with oxazolone or vehicle control, CD4^+^ T cells from Foxp3-Tocky mice were analysed. Displayed are Timer-Blue vs. Timer-Red flow cytometry plots from Vehicle or Oxazolone dLN or Ear skin. (**C**) Summary of data in **B** showing Timer-Blue and Timer-Red intensity in Timer^+^ T cells in the dLN and skin of oxazolone challenged mice, n=7-9. (**D**) Timer-locus model for analysing Timer expression. (**E**) Kernel density distributions of Timer-Angle values in dLN or skin of Oxazolone challenged Foxp3-Tocky mice. (**F**) Mean Timer-Angle values in dLN and skin of Oxazolone challenged Foxp3-Tocky mice. (**G**) Timer locus analysis of CD4^+^ T cells from the dLN (n=9) and skin (n=7) of oxazolone challenged mice. Bars represent mean +/-SD. Data are from two independent experiments.

Blue-high Red-high expression is the hallmark of high-frequency persistent transcription, which can be the most effectively identified by trigonometric data transformation to calculate the angle from the Blue axis (Timer-Angle) and the distance from the origin of individual cells (Timer-Intensity) in the Blue-Red plane (20) (**Fig. 3D**). Timer expression starts from Blue+Red-(the New locus), these New cells mature and acquire Red Fluorescence within 4 hours (**Fig. 1C and 1D**). When Timer transcription persists, cells eventually reach a balanced steady-state for Blue and Red fluorescence and accumulate in Blue+Red+ Persistent locus around 45° degree from the Blue axis. When Timer transcription is arrested, cells lose Blue fluorescence and stay in the Blue-Red+ Arrested locus while Red proteins decay (half-life is 20 h (21)). Cells in the Arrested locus can however immediately acquire Blue fluorescence again when they re-initiate *Foxp3* transcription (**Fig. 3D**), indicating that the Timer-Angle between Persistent - Arrested loci represents the recent frequency of *Foxp3* transcriptional activity (20).

The distribution (precisely, kernel-density estimation, i.e. a smoothed version of histogram) of Timer-Angle values was markedly right-skewed and high values in dLN, whether Vehicle or Oxazolone treated, while the Timer-Angle distribution in the skin showed peaks around 0° and 60° (**Fig. 3E**), with a mean Timer-Angle value of 45° (**Fig. 3F**). We classified cells according to Timer-Angle: New (Angle = 0), Persistent (Angle 30° - 60°), and Arrested (Angle = 90°). The areas between New and Persistent and between Persistent and Arrested are designated as NP-t and PA-t, respectively (20). This Timer Locus analysis clearly showed that skin-infiltrating T cells were enriched with New, NP-t and Persistent cells, compared to dLN T cells (**Fig. 3G**). Collectively these results indicate that flux from Foxp3-to Foxp3+ T cells is increased at sites of inflammation, and that skin-infiltrating Foxp3+ T cells have more active *Foxp3* transcription and are short-lived. These cells are enriched with newly born cells, and may have rapid cell divisions and/or accelerated cell death.

### Foxp3 autoregulation maintains the temporal dynamics of persistent *Foxp3* transcription

The dynamic regulation of *Foxp3* transcription during the immune response suggests that Foxp3 protein itself is involved in the regulation of *Foxp3* transcription. In order to understand the mechanism regulation of temporally-dynamic *Foxp3* transcription, we addressed this possibility by investigating the dynamics of *Foxp3* transcription in the absence of Foxp3 protein. To achieve this, we crossed Foxp3-Tocky mice with *Foxp3^Egfp/Scurfy^* mice. Treg from females from this line will all express the Foxp3-driven Timer protein, but half will express Foxp3/EGFP and the other half (EGFP-negative) express the non-functional Scurfy Foxp3 mutant protein (**Fig. 4A**) due to random X-inactivation (23, 24). This triple transgenic system thus permits the analysis of *Foxp3* transcriptional activity in the absence of functional Foxp3 protein. Foxp3-Tocky *Foxp3^Egfp/Scurfy^* female mice were immunised with Oxazolone, and subsequently, *Foxp3* expression within the dLN was analysed. All Timer+ T cells were analysed and divided into GFP+ (WT Foxp3) or GFP-(Scurfy Foxp3 mutant) expressing Treg. Interestingly, the majority of Timer+ cells were Blue+ in WT Foxp3 cells, while the majority of Timer+ cells were Blue-in Scurfy cells (**Fig. 4B**). Timer-Blue levels were remarkably decreased, and Timer-Red levels were also moderately decreased, in Scurfy cells (**Fig. 4C**). Timer-Angle was significantly higher and approached 90° in Scurfy cells (**Fig. 4D**). Scurfy Timer+ cells had very few cells displaying Persistent or PA-t transcriptional dynamics compared to GFP+ Treg, with nearly all cells located in the Arrested locus (**Fig. 4E**). Furthermore, in the absence of Foxp3 protein, Timer-expressing Persistent/PA-t cells actively transcribed *Il2* (**Fig. 4F**), a characteristic of antigen-stimulated effector T cells. These findings reveal that Foxp3 protein is itself required for controlling *Foxp3* transcription, and the Foxp3 autoregulatory loop actively operates in reactive Treg, suppressing their effector function.

**Figure 4:**
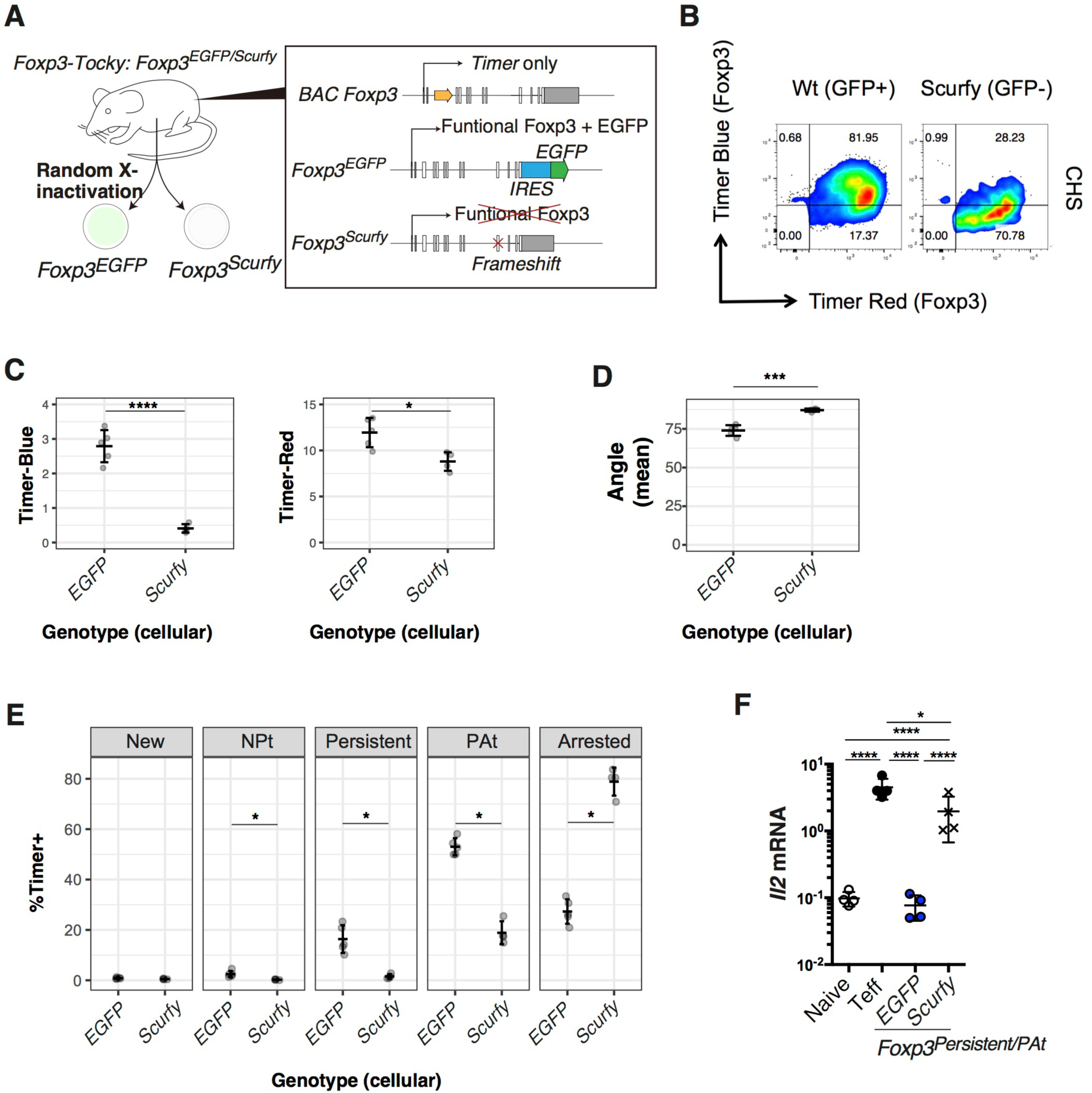
Foxp3 autoregulation maintains the temporal dynamics of persistent *Foxp3* transcription. (**A**) Design of *Foxp3-Tocky*:*Foxp3^EGFP/Scurfy^* triple transgenic system to analyse the role of functional Foxp3 in regulation of *Foxp3* transcription. *Foxp3-Tocky Foxp3^EGFP/Scurfy^* female mice were sensitized with Oxazolone for 5 d before analysis of superficial LN (sLN). (**B**) Shown are Timer-Blue vs. Timer-Red expression on Timer^+^ T cells within GFP^+^ (i.e. WT) or GFP^-^ (i.e. scurfy) T cells. (**C**) Timer-Blue and Timer-Red MFI in Timer^+^ T cells from GFP or scurfy expressing Treg. (**D**) Mean Timer-Angle values of *Foxp3^EGFP^* and *Foxp3^Scurfy^* cells, n=4. (**E**) Proportion of Timer^+^ cells within the 5 Timer loci in *Foxp3^EGFP^* nd *Foxp3^Scurfy^* from sLN (n = 4 mice). (**F**) qPCR for *Il2* mRNA in Naïve, CD44^hi^ effector T cells (Teff) and *Foxp3* Persistent cells (Blue^+^Red^+^) from GFP^+^ (*Foxp3^EGFP^*) and GFP^-^ (*Foxp3^Scurfy^*). Cells were sorted and RNA was extracted for qPCR, n=4. Bars represent mean +/-SD. Data combined from two experiments.

### Persistent *Foxp3* transcription promotes the effector Treg programme

These results suggest that persistent *Foxp3* transcription defines Treg with a unique functional status. In order to address this hypothesis, we analysed the influence of *Foxp3* transcription status on transcriptional profile using RNA-seq. T cells were isolated from mice immunised with Oxazolone, and sorted into 4 populations based on their Timer-Blue and-Red expression pattern: ‘Persistent’ *Foxp3* transcriptional expressors were isolated as Pers1 (Blue-intermediate Red-low) and Pers2 (Blue-high Red-high), in addition to PA-t (Blue-low Red-high) and Arrested (Blue-negative Red-intermediate) Foxp3 expressors (**Fig. 5A**). Each population exhibited discrete Timer-Blue,-Red, and-Angle values (**Fig. 5B-D**), confirming the successful flow sorting of the four distinct populations. Principal Component Analysis (PCA) showed that the persistent *Foxp3* expressors (i.e. Pers1 and Pers2) were clustered and similar to each other, and distinct from PA-t and Arrested *Foxp3* expressors, which made another cluster (**Fig. 5E**). Differential expression analysis identified unique clusters of genes (**Fig. 5F**). Persistent *Foxp3* expressors highly expressed the co-inhibitory/costimulatory molecules *Ctla4, Icos*, and *Tigit*, and the serine protease *Gzmb*, both of which are all involved in cancer immunity (25). In addition, persistent *Foxp3*-expressors transcribed other effector-Treg signature genes such as *Nkg7, Fgl2*, and *Irf4*, while they had low expression of naïve T cell-specific genes such as *Ccr7, Bach2*, and *Il7r*. These features are fully compatible with the previously reported effector-Treg phenotype (12, 26). The specific increase in the activation gene *Mki67* (Ki67) in persistent Foxp3 expressors also supports the idea that these cells are highly activated. Intriguingly, Pearson correlation analysis showed that *Ctla4* and effector-Treg markers (*Icos, Lag3, Tigit, Ccr4*) had relatively low correlation to Foxp3 levels (R^2^ was between 0.30 – 0.65) (**Supplementary Fig. 1A**), while these markers showed very high correlation to the activation gene, *Mki67* (R^2^ was between 0.90 – 0.98) (**Supplementary Fig. 1B**). This suggests that the T cell activation process predominantly regulates these markers and that persistent Foxp3 expressors express these markers because they are highly activated. In contrast, *Tnfrsf18* (GITR) and *Tnfrsf4* (OX40) showed high correlations to *Foxp3* (R^2^ = 0.91 and 0.82, respectively), but only moderate correlations to Mki67 (R^2^ = 0.49 and 0.74, respectively), suggesting that Foxp3 may more closely regulate GITR and OX40. *Il2ra* is ubiquitously highly expressed by all Foxp3+ cells, and not by Foxp3-cells, and showed only low correlations to both *Foxp3* and *Mki67* levels (R^2^ = 0.26 and 0.43), suggesting that CD25 expression is stabilized in Foxp3+ cells but regulated primarily by other factors, such as IL-2 signalling once Foxp3 is expressed (**Supplementary Fig. 1A and 1B**). Persistent Foxp3 expressor-specific genes were enriched with cell cycle-related pathways. In contrast, the PA-t and Arrested-specific genes were enriched with cytokine-signalling pathways but not cell-cycle-related pathways (**Fig. 5G**). Collectively, RNA-seq analyses showed that temporally-persistent *Foxp3* transcription is an underlying mechanism for effector Treg differentiation.

**Figure 5:**
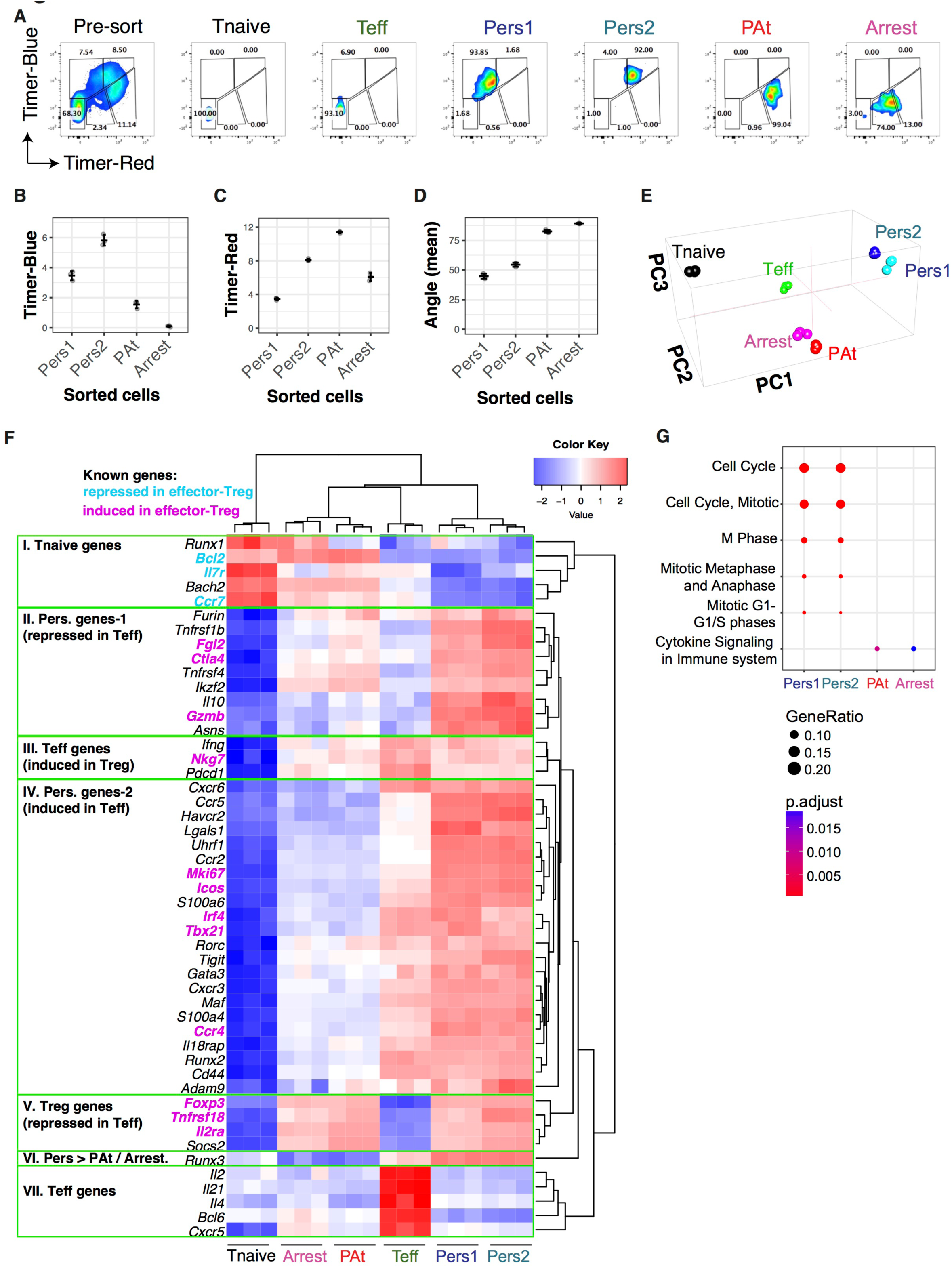
Persistent *Foxp3* transcription promotes the effector Treg programme. Foxp3-Tocky mice were sensitised with Oxazolone for 5 days and dLN harvested and stained with CD4 and CD44 for sorting. (**A**) Pre and post-sort purities displaying Timer-Blue vs Timer-Red fluorescence for Tnaive (CD4^+^CD44^lo^Timer^-^), Teff (CD4^+^CD44^hi^Timer^-^). Timer^+^ T cells from sensitised *Foxp3^Timer^* mice were sorted into Pers1, Pers2 PA-t and Arrested T cell subsets, according to the maturation of Timer chromophore n = 3 groups, each made from pools of 3 mice. Mean Timer-Blue (**B**) or Timer-Red fluorescence intensity levels in the sorted Timer+ subsets, n=3. (**D**) Timer Angle values for sorted Timer^+^ T cell subsets. RNA was extracted from the samples and sequencing was performed as described in the methods. (**E**) Cluster analysis of the transcriptomes of the sorted samples by PCA. (**F**) Heatmap analysis of differentially expressed genes (DEGs) in the 4 Timer^+^ subsets, compared to Tnaive and Teff, grouping genes into 7 categories. (**G**) Pathway analysis of the 4 DEG gene lists.

### Identification of distinct groupings of cell surface markers for targeting T cells with specific *Foxp3* transcriptional dynamics

Next, we analysed protein expression of candidate surface proteins identified by RNA-seq for targeting T cells with specific *Foxp3* transcriptional dynamics in vivo. T cells from Foxp3-Tocky mice were immunized with Oxazolone, and analysed by flow cytometry for the expression levels of candidate markers from Fig. 5 in each Timer Locus. Hierarchical clustering of MFI levels showed that these candidate genes were grouped into two main groups, in addition to Neuropilin 1 (Nrp1) which had its own separate cluster (**Fig. 6A**). Group I markers, showed high expression throughout New – Persistent *Foxp3* expressors, but fell drastically as Blue fluorescence decreases in PA-t and Arrested *Foxp3* expressors (**Fig. 6B**), suggesting that these markers are associated with the initiation of new *Foxp3* transcription in activated T cells and with high-frequency *Foxp3* transcription. Group II markers are less expressed in new *Foxp3* expressors, and increased in persistent and PA-t *Foxp3* expressors (**Fig. 6C**), suggesting that these markers can be sustained by moderate frequencies of *Foxp3* transcription once the Foxp3 autoregulatory loop is established. Nrp1 is characteristically high in PA-t *Foxp3* expressors, which may explain why this marker has been used to identify thymus-derived Treg (**Fig. 6D**). Importantly, the expression levels of both Group I and II markers markedly declined in Arrested *Foxp3* expressors. Together these results further support the notion that the temporal frequency (rate) of *Foxp3* transcription regulates the Treg phenotype and function.

**Figure 6:**
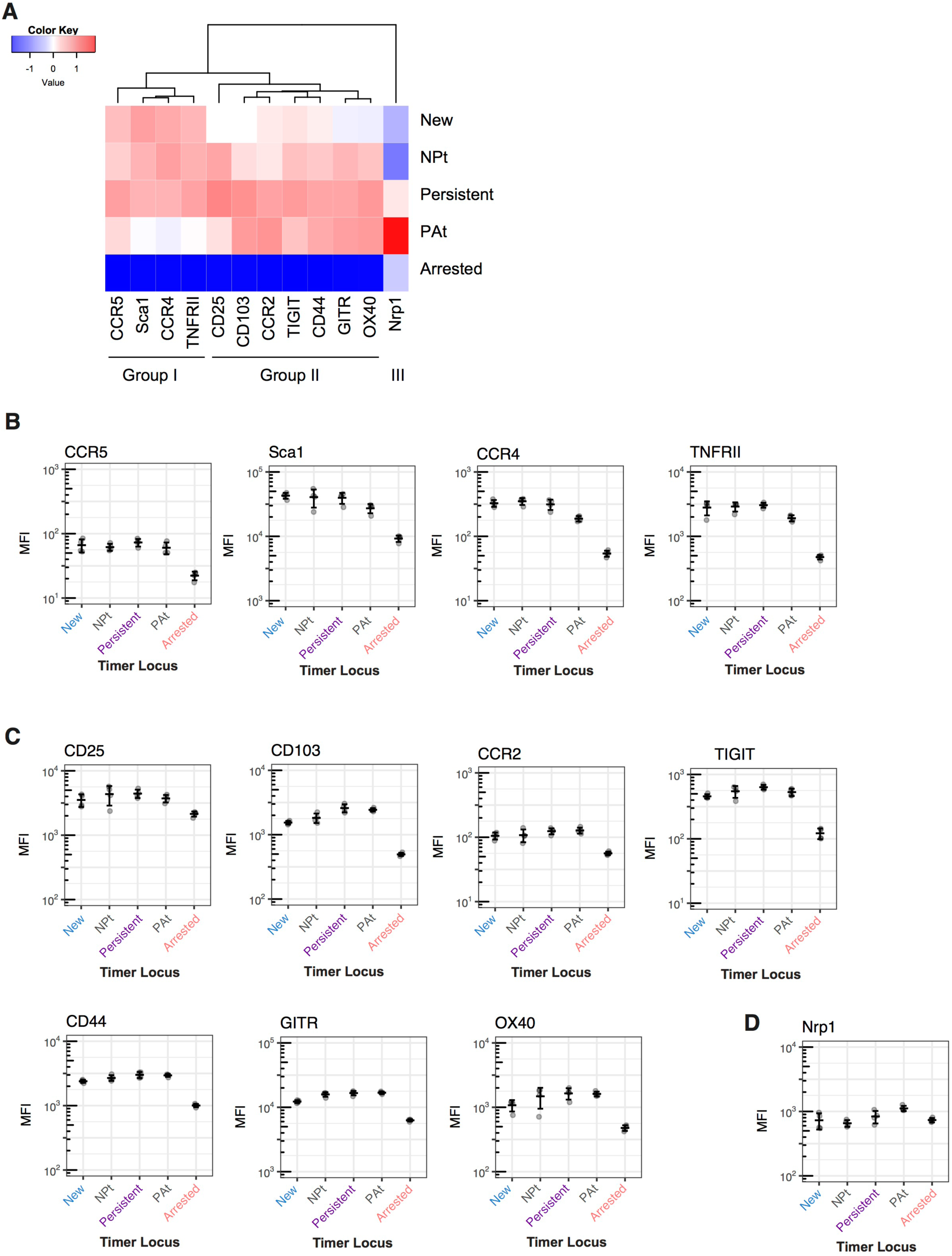
Identification of distinct groupings of cell surface markers for targeting T cells with specific *Foxp3* transcriptional dynamics. Foxp3-Tocky mice were sensitized with Oxazolone for 4 days, and then dLN were harvested and CD4+ T cells analysed for Timer expression and key membrane proteins identified as differentially expressed by RNA-seq. (**A**) Heatmap analysis of the expression of different surface proteins according to cells within different Timer loci. These findings generate 3 main groups. (**B**) MFI expression levels of Group I, (**C**) group II or (**D**) group III surface markers in relation to Timer-loci of the cells, n=4. Bars represent mean +/-SD.

### Foxp3-Tocky allows visualistion of the manipulation of T cells with specific *Foxp3* transcriptional dynamics upon immunotherapy

We asked if surface proteins from Group I and Group II in Fig. 6 can be used to target T cells with specific *Foxp3* transcriptional dynamics. If successful, this approach may lead to future precision immunotherapy for controlling Foxp3-mediated immune regulation by manipulating its temporal dynamics. Both TNFRII (Group I) and OX40 (Group II) are targets for immunotherapy and may be used to manipulate T cell responses, and currently clinical trials are ongoing for melanoma and other solid cancers (27). However, it is unknown whether these immunotherapy antibodies affect *Foxp3* transcriptional dynamics.

We used the CHS model, and antibodies were administered during the challenge phase (**Fig. 7A**). Treatment with anti-TNFRII antibody increased the frequency of Blue+Red-T cells, and Blue+Red+ T cells at the inflamed site (**Fig. 7B)**. Although average Blue and Red levels were overall not affected by the antibody (**Fig. 7C and 7D**), Timer-Angle tended to be lower within the anti-TNFRII treated group (p=0.055), likely reflecting the increased proportion of pure blue T cells at the inflamed site (**Fig. 7E**). Analysis of the Timer-Angle distribution revealed that Isotype control treatment showed one dominant peak around the 45° mark, whilst anti-TNFRII treatment produced a more flattened, peak from 25-50 degrees (**Fig. 7F**). Analysis of Timer locus revealed that anti-TNFRII increased both New and NPt stage *Foxp3* expressors, indicating that anti-TNFRII immunotherapy enhances the flux of Foxp3-into Foxp3+ T cells (**Fig. 7G**). There was no significant change in the high-frequency/persistent *Foxp3* transcribers, and anti-TNFRII treatment did not significantly alter the course of inflammation (**Fig. 7H**). In contrast, anti-OX40 treatment appeared to dramatically reduce the frequency of Blue+Red+ Foxp3+ T cells in inflamed skin (**Fig. 7I**). Whilst Blue levels were unaffected (**Fig. 7J**), both Timer-Red and Timer-Angle were significantly lower in anti-OX40 treated mice (**Fig. 7K and 7L**). Interestingly, the peak of Timer-Angle distribution in anti-OX40 treated mice was shifted towards the NPt (**Fig. 7M**). Timer locus analysis revealed that anti-OX40 reduced the frequency of more mature *Foxp3* expressors, as both Persistent and PA-t loci were significantly reduced in anti-OX40 treated mice (**Fig. 7N**) These data indicate that, in the presence of anti-OX40 antibody, Treg are very short lived at the inflamed site, and that high-frequency *Foxp3* expressors are either depleted or undergo activation-induced death. Interestingly, this reduction in high-frequency *Foxp3* expressors led to delayed resolution of inflammation (**Fig. 7O**), demonstrating the importance of sustained *Foxp3* transcription for regulating the T-cell response. In summary, both of the effects of the immunotherapies tested were in keeping with the *Foxp3* transcriptional dynamics identified in Figure 6, showing how Foxp3-Tocky may be used to identify new therapeutic targets and mechanism of drug action in pre-clinical models.

**Figure 7:**
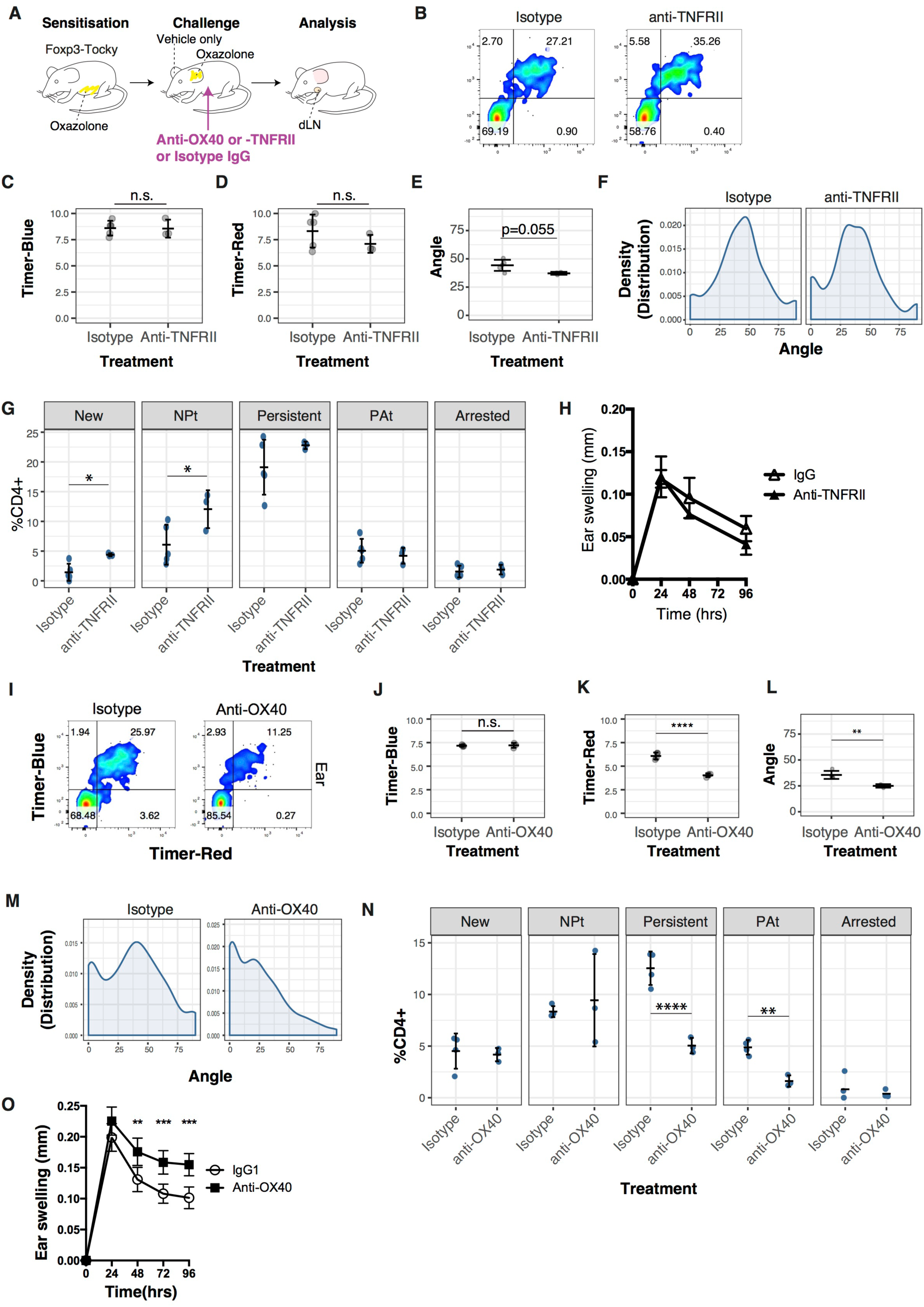
Foxp3-Tocky allows visualistion of the manipulation of T cells with specific *Foxp3* transcriptional dynamics upon immunotherapy. (**A**) Experimental design for the administration of anti-OX40 or anti-TNFRII antibody to target the challenge phase of CHS. (**B**) Mice were challenged with Oxazolone and then administered Isotype or anti-TNFRII antibodies. 96hrs later, CD4+ T cells from the skin were analysed for expression of Timer-Blue vs. Timer-Red. Summary of Timer-Blue MFI (**C**) or Timer-Red MFI (**D**) or Timer-Angle (**E**) in skin infiltrating Timer+ T cells, Isotype n=5, anti-TNFRII n=3. (**F**) Kernel-density distributions of Timer+ T cells in Isotype or anti-TNFRII treated mice 96 h after challenge with Oxazolone. (**G**) Timer-locus distribution of skin infiltrating CD4+ T cells in Isotype or anti-TNFRII treated Foxp3-Tocky mice. (**H**) Ear swelling during the challenge phase of CHS in Isotype or anti-TNFRII treated Foxp3-Tocky mice, bars represent mean +/-SEM. (**I**) Mice were challenged with Oxazolone and then administered Isotype or anti-OX40 antibodies. 96hrs later, CD4+ T cells from the skin were analysed for expression of Timer-Blue vs. Timer-Red. Summary of Timer-Blue MFI (**J**) or Timer-Red MFI (**K**) or Timer-Angle (**L**) in skin infiltrating Timer+ T cells, Isotype n=4, anti-OX40 n=3. (**M**) Kernel-density distributions of Timer+ T cells in Isotype or anti-OX40 treated mice 96 h after challenge with Oxazolone. (**N**) Timer-locus distribution of skin infiltrating CD4+ T cells in Isotype or anti-OX40 treated Foxp3-Tocky mice. (**O**) Ear swelling during the challenge phase of CHS in Isotype (n=19) or anti-OX40 (n=18) treated mice. Data in (**O**) are combined from two experiments and show mean +/- 95% confidence intervals. Unless stated, all other graphs show mean +/-SD.

## Discussion

Sustained *Foxp3* transcription is a cardinal feature of the functional maturation of Treg, or effector-Treg (**Fig. 5**). Importantly, sustained *Foxp3* transcription requires functional Foxp3 protein itself. This type of transcriptional regulation (i.e. transcription factor regulates its own gene) is defined as positive autoregulation, which allows a slow and sustained transcriptional response to external signalling cues (28). This may constitute the mechanism behind effector Treg differentiation in inflammatory environments. It is known that, among the conserved non-coding DNA sequence (CNS) elements of the *Foxp3* gene, the CNS2, which contains the TSDR, is required for maintenance of Foxp3 expression after cell division (29). The CNS2 regions is bound by Foxp3 (29), Runx1/Cbf-|β (30, 31), and Stat5 (32). Thus, the autoregulatory transcriptional circuit of Foxp3 may involve the Foxp3-Runx1 interaction (5) at the CNS2 and IL-2 signalling. The interaction of Foxp3 and Runx1 is critically important for the repression of *Il2* and *Ifng* (5). In agreement with this, functional Foxp3 protein was required for Persistent/ PA-t cells, in which the autoregulatory loop is established, to suppress *Il2* and *Ifng* transcription. Furthermore, this autoregulation may be essential for establishing and consolidating the effector Treg programme. Given that the autoregulatory transcriptional circuit sustains *Foxp3* transcription, genes with linear correlations to *Foxp3* transcripts (e.g. *Tnfrsf4* and *Tnfrsf18*, Supplementary Fig S1) are likely regulated by this transcriptional circuit also. On the other hand, genes with high expression in persistent *Foxp3* expressors that show moderate correlations to *Foxp3* levels (e.g. *Ctla4*, Supplementary Fig S1A) are likely to be regulated by both the T cell activation process and Foxp3-autoregulation dependent mechanisms. Thus, in activated effector Treg, the autoregulatory loop of the *Foxp3* gene and the T cell activation process may cooperatively control transcellular suppressive mechanisms (e.g. CTLA-4), while suppressing the effector functions (e.g. cytokine production) in a cell-intrinsic manner. A future study may focus on the identification of molecular mechanisms for the establishment and control of the Foxp3 autoregulatory transcriptional circuit across chromatin regions and/or at the single cell level.

Given that most activated T cells are destined to die after resolution of inflammation (33), the majority of new and persistent *Foxp3* expressors in tissues should also die. It is plausible that Foxp3 may drive apoptosis of these activated T cells after the resolution of inflammation, since Foxp3 has been demonstrated to be pro-apoptotic in the absence of survival signals (i.e. cytokine-deprivation death) (34). If some of the reactive Foxp3 expressors survive beyond the resolution of inflammation, such cells will be memory T cells by definition. Some cells may sustain Foxp3 expression and survive as “memory Treg”, and some others may lose Foxp3 expression after the resolution of inflammation and survive as memory-phenotype (memory-like) T cells (16). Intriguingly, ex-Treg are enriched in the memory T cell pool and prone to express Foxp3 more readily than naïve T cells (14, 35). These hypotheses will be best addressed in future studies of T cell memory, and will require new approaches to combine the Tocky technology and the analysis of cellular dynamics several weeks after the onset of inflammation.

In the hapten model, ~10% of Timer+ cells are identified at the New locus in the skin (Fig. 3G). Given that these new cells quickly move to NP-t/Persistent locus within 4 h (**Fig. 1B**), a significant proportion of cells at the Persistent locus may have been supplied from new cells within 72 h before the analysis. It is also possible that cells at the PA-t and Arrested loci move to the Persistent locus by increasing *Foxp3* transcription. These two possibilities are not mutually exclusive and are compatible with evidence that the effector Treg population arises from both thymic and peripherally-induced Treg (12), while the ratio of the contributions may vary between antigens (16). Future studies using TCR repertoire sequencing will fully answer this question.

The current study has shown that Foxp3-Tocky is effective in designing immunotherapy strategies to target a specific phase of *Foxp3* transcriptional dynamics, and also, for visualising alterations of the dynamics upon immunotherapy. In fact, we successfully manipulated in vivo *Foxp3* dynamics by targeting Timer-stage specific surface receptors. By targeting Group I (TNFRII) or Group II (OX40) marker of *Foxp3* dynamics (**Fig. 6**) we either increased the flux of new *Foxp3* transcription (anti-TNFRII) or decreased persistent *Foxp3* expressors (anti-OX40). While the precise cellular mechanism is to be determined, anti-OX40 treatment resulted in either the depletion or death of persistent *Foxp3* expressors within the skin, which are enriched with effector Treg. OX40 is an emerging target for cancer immunotherapy, and its agonistic antibodies for augmenting the effector function of T cells are currently tested in clinical trials (36). It is currently thought that OX40 is expressed mainly by activated effector T cells, and is also constitutively expressed on all Treg (37). Such OX40 high expression is found relatively more frequently on persistent Foxp3 expressors compared to Foxp3-negative effector T cells.

In summary, we have dissected the previously hidden dynamics of *Foxp3* transcription that regulate the functions of Treg through time. In addition, we have shed light on how Foxp3-mediated molecular mechanisms coordinate T cell responses in existing Treg and non-Treg, introducing a new dimension into studies on Foxp3-driven T cell regulation.

## Methods

### Study Design

Sample size for anti-OX40 CHS experiments was estimated based on a power calculation with alpha= 0.05 and power 80%, which gave a sample size of 8. For analysis of *Foxp3-Tocky Foxp3*^*EGFP/Scurfy*^ mice, 4 mice were analysed over two independent experiments (age 6-10 weeks). For anti-OX40 experiments, a pilot experiment was performed to obtain estimates for effect size and standard deviation, which yielded an experimental group size of 9. Two independent experiments were then performed to test the null hypothesis that anti-OX40 treatment had no effect on ear swelling at 96 h. The null hypothesis was rejected in both experiments, with no significant inter-experiment variation. Therefore, data from both experiments were pooled for presentation of results. For all other experiments a minimum of 3 animals were used. Where male and female mice were used in the same study, these were randomised into experimental groups according to sex and age. Similarly, wild type littermates from transgenic mice were randomised into experimental groups for the anti-OX40 experiments. The individual making the ear measurement was blinded to the treatment group of the mouse until after the measurement was taken.

### Transgenesis and mice

*Foxp3*^-^Tocky and Foxp3-Tocky:Foxp3-IRES-GFP transgenic reporter strains were generated as described (20). The triple transgenic *Foxp3-Tocky Foxp3*^*EGFP/Scurfy*^ mice were generated by crossing *Foxp3-Tocky* mice with *Foxp3^tm9(EGFP/cre/ERT2)Ayr^/J* and *B6.Cg-Foxp3*^*sf*^/J (Jackson Laboratories, #016961 and #004088, respectively). All animal experiments were performed in accordance with local Animal Welfare and Ethical Review Body at Imperial College London (Imperial) and University College London (UCL), and all gene recombination experiments were performed under the risk assessment that was approved by the review board at Imperial and UCL.

### In vitro cultures for determining mRNA half lives

CD4+ T cells from the spleens of Foxp3-Tocky mice were isolated by immunomagnetic separation (StemCell technologies) and cultured at 4x10^5^ cells per well in the presence of 10μg/ml Actinomycin D. Cells were isolated at the indicated time points, and RNA was extracted, and cDNA synthesises as described below.

### In vitro cultures for determining Timer-Blue half-life

Naïve T cells from *Foxp3-Tocky* mice were isolated by negative selection using immunomagnetic selection (StemCell Technologies) and 2x10^5^ cells cultured on anti-CD3 (clone 1452C11, 2 μg/ml) and anti-CD28 (clone 37.51,10μg/ml; both eBioscience)-coated 96 well plates (Corning) in the presence of 500 U/ml rhIL-2 (Roche) and 5 ng/ml rhTGFβ (R&D) for 48 hrs in a final volume of 200 μL RPMI1640 (Sigma) containing 10% FCS and penicillin/streptomycin (Life Technologies). Cells were harvested then incubated with 100μg/ml cycloheximide (Sigma) for the indicated time points before analysis by flow cytometry.

### Contact Hypersensitivity and antibody treatments

Oxazolone (Sigma) solutions were prepared fresh in ethanol for each experiment. For sensitisation, 150 μL of 3% Oxazolone was applied to the shaven abdominal skin of anaesthetised 5-10 week old mice. Five days later, left and right ear thickness were measured, and the elicitation phase started by the application of 20 μL of a 1% Oxazolone solution to one ear, or 20 μL of vehicle (100% ethanol) to the contralateral ear. Twenty-four to ninety-six h later, ear thickness was measured using a digital micrometer (Mitutoyo). In anti-OX40 experiments, 0.5 mg of anti-OX40 (OX86, BioXCell) or rat IgG1 (MAC221, kind gift from Prof. Anne Cooke) were injected i.p. on day of challenge and 48 h later. For anti-TNFRII experiments, 0.5 mg of anti-TNFRII (TR75-54.7 BioXcell) or Armenian hamster IgG (BioXcell) were injected intra-peritoneally (i.p). on day of challenge.

### Flow cytometric analysis and cell sorting

Following spleen or thymus removal, organs were forced through a 70 μm cell strainer to generate a single cell suspension. For splenocyte preparations a RBC-lysis stage was employed. Staining was performed on a V-bottom 96-well plate, or in 15 ml falcon tubes for cell sorting. Analysis was performed on a BD Fortessa III instrument. The blue form of the Timer protein was detected in the blue (450/40 nm) channel excited off the 405 nm laser. The red form of Timer protein was detected in the mCherry (610/20) channel excited off the 561nm laser. For all experiments a fixable eFluor 780-fluorescent viability dye was used (eBioscience). The following directly conjugated antibodies were used in these experiments: CD4 APC (clone RM4-5, eBioscience), CD4 Alexa-fluor 700, CD4 BUV395 (clone GK1.5 BD Biosciences), (clone RM4-5, Biolegend), CD8 PE-Cy7 (clone 53-6.7, Biolegend), CD8 BUV737 (clone 53-6.7, BD Biosciences) TCRβ FITC & Alexafluor 700 (clone H57-597, Biolegend), TCRβ BUV737 (clone H57-597, BD Biosciences), CD25 PerCPcy5.5 (PC61.5, eBioscience) or PE-Cy7 (PC61.5, Tombo Bioscience), CD44 APC (clone IM7, eBioscience) or Alexafluor 700 (clone IM7, Biolegend), OX40 PE-Cy7 and APC (clone OX86, Biolegend), GITR FITC (clone DTA-1, Biolegend), TNFRII APC (clone TR7554, R&D Systems), Neuropilin-1 APC or PE-Cy7 (clone 3E12 Biolegend), CD103 APC (clone 2E7, Biolegend) CCR4 APC (clone 2G12, Biolegend), TIGIT PE-Cy7 (Clone 1G9, Biolegend), CCR2 AF647 (clone SA203G11, Biolegend), CCR5 APC (clone HM-CCR5, Biolegend and Sca1 (clone D7, Biolegend).

### qPCR analysis

For sorted cells, RNA was extracted using the Arcturus PicoPure RNA kit (Life Technologies) according to the manufacturer’s instructions. cDNA was generated using random hexamers and Superscript II (Life Technologies) according to the manufacturer’s instructions. mRNA expression was quantified using SYBR green (BioRad) and expressed relative to the housekeeping gene *Hprt*. Primer sequences: *Hprt* For: AGCCTAAGATGAGCGCAAGT, *Hprt* Rev: TTACTAGGCAGATGGCCACA. *Il2* For: AGCAGCTGTTGATGGACCTA *Il2* Rev: CGCAGAGGTCCAAGTTCAT (38). *Foxp3* For: CACCCAAGGGCTCAGAACTTCTAG, *Foxp3* Rev: ATGACTAGGGGCACTGTAGGCA, *Foxp3-Timer* For: CAGCTCCTCTGCCGTTATCC, *Foxp3-Timer* Rev: CCTCGCCCTCGATCTCGA

### RNA-seq

*Foxp3-Tocky* mice were sensitised by application of 3% oxazolone to the abdomen. Five days later, dLN were harvested and CD44^lo^Foxp3^-^, CD44^hi^Foxp3^-^, *Pers1, Pers2, PAt*, and *Arrested* T cells were sorted and RNA extracted using the Arcturus PicoPure RNA kit (Life Technologies) according to the manufacturer’s instructions. Library was prepared using the Illumina TruSeq^®^ Stranded mRNA LT kit according to manufacturer’s instructions. Libraries were analysed by Bioanalyzer (Agilent) for quality control and quantified using Qubit (Life Technologies). Library concentrations were normalised and pooled and sequenced on the Illumina HiSeq platform (Illumina) and 100 bp paired-end readings were obtained. After demultiplexing fastq files and performing QC analysis by *fastQC*, sequence reads were aligned to either the mouse genome (mm10) with or without the *Timer* gene by *TopHat2*. Statistical analysis was performed using the Bioconductor package *DESeq2*. Differentially expressed gene lists were generated using the union of the genes with FDR < 0.05 and log fold change > 1 in comparison to any of the other 3 sample groups (i.e. Pers1 vs PA-t, Arrested, and CD44^hi^). PCA was performed using the CRAN package *Stats*, and the 3D-plot was generated by the CRAN package *rgl*. Heatmap was generated by the CRAN package *gplots*, and hierarchical clustering used a complete linkage algorithm. Pathway analysis was done using the Bioconductor package *clusterProfiler* and the Reactome database through the package *ReactomePA*.

### Timer data analysis

Sample data including a negative control were batch gated for T cell populations and exported by FlowJo (FlowJo LLC, OR) into csv files, including all compensated fluorescence data in the fcs file. The code developed in this study imports csv files into R, preprocesses and normalises data, and automatically identifies Timer^+^ cells and performs Trigonometric data transformation, producing Timer-Angle and-Intensity data for individual cells in each sample, as previously described (20). Basic procedures for flow cytometric data analysis have been previously described elsewhere (19).

### Statistical analysis and data visualisation

Statistical analysis was performed on R or Prism 6 (Graphpad) software. Percentage data for Timer^+^ and Timer locus analysis was analysed by Mann-Whitney U test or Kruskal-Wallis test with Dunn’s multiple comparisons using the CRAN package *PMCMR*. Samples with fewer than 20 Timer^+^ cells were not included in the analysis (except for Fig. 7G, where threshold was set as 10). For analysis of mRNA, ear thickness data, and Timer-Angle and Intensity, Student’s t-test was used for comparison of two means. For comparison of more than two means, a one-way ANOVA with Tukey’s post-hoc test was applied using the CRAN package *Stats*. For comparison of variation between two data sets across two variables, a two-way ANOVA was used with Sidak’s multiple comparisons test. Scatterplots and density plots were produced by the CRAN packages *ggplot2* and *graphics*. All computations were performed on Mac (version 10.11.6). Adobe Illustrator (CS5) was used for compiling figures and designing schematic figures. Variance is reported as SD or SEM unless otherwise stated. * p<0.05, ** p<0.01, *** p<0.001, **** p<0.0001.

## Data and code availability

RNA-seq data can be obtained at NCBI GEO with the accession number GSE89481. All R codes are available upon request. Data will be made available upon reasonable requests to the corresponding author.

## Author Contributions

M.O. conceived the Tocky strategies and the project. D.B., T.C, and M.O conceived and designed immunological experiments. D.B. and P.P.M performed animal experiments. D.B. and A.P. performed molecular experiments. D.B. performed RNA-seq experiment. M.O. wrote computational codes, and performed bioinformatics analysis and data visualisation. D.B., T.C., and M.O. wrote the manuscript.

## Acknowledgements

We thank for their kind support at the Flow Cytometry facility, Dr. Ayad Eddaoudi and Ms Stephanie Canning (University College London), and also Ms Jane Srivastava or Ms Catherine Simpson (Imperial College London). M.O. is a David Phillips Fellow (BB/J013951/2) from the Biotechnology and Biological Sciences Research Council (BBSRC). T.C. is supported by Great Ormond Street Hospital Children’s Charity and the National Institute of Health Biomedical Research Centre at Great Ormond Street Hospital for Children NHS Foundation Trust and University College London. We thank Dr. Miho Ishida (University College London) for technical advice and help.

